# Dynamic evolutionary history and gene content of sex chromosomes across diverse songbirds

**DOI:** 10.1101/454843

**Authors:** Luo-hao Xu, Gabriel Auer, Valentina Peona, Alexander Suh, Yuan Deng, Shao-hong Feng, Guo-jie Zhang, Mozes P.K. Blom, Les Christidis, Stefan Prost, Martin Irestedt, Qi Zhou

## Abstract

Songbirds have a species number almost equivalent to that of mammals, and are classic models for studying mechanisms of speciation and sexual selection. Sex chromosomes are hotspots of both processes, yet their evolutionary history in songbirds remains unclear. To elucidate that, we characterize female genomes of 11 songbird species having ZW sex chromosomes, with 5 genomes of bird-of-paradise species newly produced in this work. We conclude that songbird sex chromosomes have undergone at least four steps of recombination suppression before their species radiation, producing a gradient pattern of pairwise sequence divergence termed ‘evolutionary strata’. Interestingly, the latest stratum probably emerged due to a songbird-specific burst of retrotransposon CR1-E1 elements at its boundary, or chromosome inversion on the W chromosome. The formation of evolutionary strata has reshaped the genomic architecture of both sex chromosomes. We find stepwise variations of Z-linked inversions, repeat and GC contents, as well as W-linked gene loss rate that are associated with the age of strata. Over 30 W-linked genes have been preserved for their essential functions, indicated by their higher and broader expression of orthologs in lizard than those of other sex-linked genes. We also find a different degree of accelerated evolution of Z-linked genes vs. autosomal genes among different species, potentially reflecting their diversified intensity of sexual selection. Our results uncover the dynamic evolutionary history of songbird sex chromosomes, and provide novel insights into the mechanisms of recombination suppression.

## Introduction

Songbirds (Oscines, suborder Passeri) have over 5000 species and comprise the majority of passerines and nearly half of the all extant bird species^1^. This is a result of the largest avian species radiation occurred about 60 million years (MY) ago^2^. Facilitated by the development of genomics, many species besides the zebra finch (*Taeniopygia guttata*) are now transforming into important models for studying molecular patterns and mechanisms of speciation^3^^,^^4^, supergenes^5^ and cognition^6^, out of their long history of ecological or behavioral studies, out of their long history of ecological or behavioral studies. One major reason that has been fueling biologists’ fascination with songbirds is their staggering and diversified sexual traits. Many species possess striking plumage forms and colors, sophisticated songs and mating rituals, all of which can undergo rapid turnovers even between sister species. Theories predict that sex chromosomes play a disproportionately large role in speciation (the ‘large X/Z’ effect), sexual selection and evolution of sexually dimorphic traits^7^^-^^9^. However, the evolutionary history of songbird sex chromosome remains unclear, because there were few genomic studies characterizing songbird sex chromosomes except for the Collared Flycatcher (*Ficedula albicollis*)^10^. In contrast to the mammalian XY system, birds have independently evolved a pair of female heterogametic sex chromosomes that are usually heteromorphic in females (ZW) and homomorphic in males (ZZ). A recent cytological investigation of over 400 passerine species found a higher fixation rate of chromosome inversions on the Z chromosome than autosomes within species. Gene flow in the Z chromosome is thus more likely reduced in the face of hybridization^11^. Indeed, a significantly lower level of introgression, and a higher level of *F*st in Z-linked genes compared to autosomal genes has been reported from studying pairs of recently diverged songbird species^12^^-^^15^. Such a large-Z pattern is probably caused by several factors which act in an opposite manner to the XY sex system. First, Z chromosomes are more often transmitted in males, thus are expected to have a higher mutation rate than the rest of the genome, due to the ‘male-driven evolution’ effect^16^. Second, as sexual selection more frequently targets males, the variation in male reproductive success will further reduce the effective population size of Z chromosome from three quarters of that of autosomes^17^. The consequential stronger effect of genetic drift is expected to fix excessive slightly deleterious mutations on the Z chromosome, and lead to a faster evolutionary rate than on autosomes (the ‘fast-Z’ effect)^18^. This has been demonstrated in the Galloanserae (e.g., chicken and duck) species, those of which undergo strong sperm competition, i.e., more intensive male sexual selection, exhibit a larger difference between the Z chromosome and autosomes in their evolutionary rates^19^.

In contrast to the avian Z chromosome, or more broadly the mammalian XY chromosomes, the genomic studies of avian W chromosomes, especially those of songbirds have not started only until recently^10^^,^^20^^,^^21^. This is because most genomic projects prefer to choose the homogametic sex (e.g., male birds or female mammals) for sequencing, in order to avoid the presumably gene-poor and highly repetitive Y or W chromosomes. The Y/W chromosomes have undergone suppression of recombination to prevent the sex-determining gene or sexually antagonistic genes (beneficial to one sex but detrimental to the other) from being transmitted to the opposite sex^22^. As a result, interference between linked loci (‘Hill-Robertson’ effect) reduces the efficacy of natural selection and drives the ultimate genetic decay of non-recombining regions of Y/W chromosomes^23^. This process can be accelerated by positive selection targeting, for example, male-related genes on the Y chromosome^24^; or by background selection purging the deleterious mutations from highly dosage-sensitive genes^25^. Simulation showed that both forces play a different role at different stages of Y/W degeneration^26^. Both have been implicated in analyses of mammalian^24^^,^^27^ and *Drosophila*^28,29^ Y-linked genes. However, no evidence has been found for female-specific selection among the W-linked genes (also called gametologs) of chicken^21^ or flycatcher^30^.

Intriguingly, in both birds^20^ and mammals^31^, as well as several plant species (e.g. *Silene latifolia*^32^), recombination suppression has proceeded in a stepwise manner presumably through chromosome inversions, leaving a stratified pattern of sequence divergence between sex chromosomRef28es termed ‘evolutionary strata’^33^. Eutherian mammalian X and Y chromosomes have been inferred to share at least three strata, with another two more recent ones shared only among catarrhines (old world monkeys and great apes)^27^. It has been recently discovered that the history and tempo of avian sex chromosome evolution is much more complicated than that of mammals^20^. All bird sex chromosomes only share the first step of recombination suppression (stratum 0, Aves S0) encompassing the avian male-determining gene *DMRT1*. This was followed by the independent formation of S1 in the Palaeognathae (e.g., ratites and tinamous) and in the ancestor of the Neognathae (all other extant avian radiations). Ratites have halted any further recombination loss and maintained over two thirds of the entire sex chromosome pair as the exceptionally long recombining pseudoautosomal regions (PAR). Therefore, their W chromosomes are unusually homomorphic and gene-rich comparing to the Z chromosomes. In contrast, all species of Neognathae examined have suppressed recombination throughout most regions of the sex chromosomes with short and varying sizes of PAR^34^. Overall, avian W chromosomes seem to have retained more genes and decayed at a slower rate than the mammalian Y chromosomes. Furthermore, sexually monomorphic species (e.g., most ratites) seem to differentiate even slower than sexually dimorphic species (chicken and most Neoaves) in their sex chromosomes, consistent with the hypothesis that sexually antagonistic genes have triggered the expansion of recombination suppression between sex chromosomes^35^. However, due to the ratites’ deep divergence from other birds, and also an expected much lower mutation rate due to their larger body size and longer generation time, it is unclear what the actual influence of sexual selection is on the rate of sex chromosome evolution. All Neoaves species share one stratum S2, with the more recent evolutionary history of sex chromosomes of songbirds unclear. So far, only one songbird, the collared flycatcher has been extensively characterized for its W-linked genes^30^, whose number is within the range of 46 to 90 W-linked genes reported for other Neoaves^20^. To elucidate the evolutionary history of songbird sex chromosomes, we produced high-quality female genomes of five birds-of-paradise (BOP). Together with a re-analysis of 6 other published female genomes of songbird species^30^^,^^36^^-^^39^, our analyses cover the two major songbird lineages (Corvida and Passerida) that rather diverged in the last 50 MY^2^^,^^40^.

## Results

### Characterization of songbird sex chromosome sequences

We produced between 36-150 fold genomic coverage of sequencing data for each BOP species, and performed *de nov*o genome assembly followed by chromosome mapping using the great tit (*Parus major*) genome as reference^6^. The high continuity and completeness of the draft genomes are reflected by their scaffold N50 lengths (all >3 Mb except for the raggiana BOP, *Paradisaea raggiana*) and BUSCO core gene completeness scores (92.9% to 94.0%) (**Supplementary Table 1**). To reconstruct the evolutionary history of sampled songbird sex chromosomes, we first identified the sex-linked sequences from the draft genome of each species. We searched for the scaffolds from sexually differentiated regions (SDR) as those that show half the female sequencing depth of autosomes (**Fig. 1A**, **Supplementary Fig. 1**). Scaffolds from putative PARs were inferred by their homology to that of flycatcher^41^, and also sequencing depth similar to autosomes. It is noteworthy that our method cannot identify very recent fusion/translocation of autosomal fragments to the sex chromosome pair (‘neo-sex’ chromosome), as in the case of warblers^42^. All the studied songbirds have a short putative PAR ranging from 564 kb to 781 kb. We then categorized SDR sequences as either Z- or W-linked with the expectation that the W chromosome would diverge much faster than the homologous Z chromosome of the same species, when being compared to the Z chromosome of an outgroup species (**Materials and Methods**), as a result of rapid accumulation of deleterious mutations and repetitive elements after recombination suppression. It is possible that Z-linked paralogs, although not expected to be abundant in the compact avian genomes^36^, would confound our identified W-linked sequences by showing a similar sequence divergence pattern. Thus, among the five species with male sequencing data, we further verified the sex-linkage by confirming all the putative W-linked scaffolds with a clear female-specific pattern (**Fig. 1A, Supplementary Fig. 2**). The assembled lengths of the largely euchromatic parts of W chromosomes range from 1.33 to 6.52 Mb, corresponding to only 1.9% to 8.5% of the Z chromosome length across species (**Fig. 1B**, **Supplementary Table 2**), probably as a result of large deletions and massive invasions of repetitive elements which might be difficult to assemble with existing technologies. Indeed, the repeat content of the assembled W chromosomes is 2.5 to 4.9 fold higher than that of Z chromosomes on the chromosome-wide average (**Supplementary Fig. 3 and Table 2**), consistent with the patterns of W chromosomes of chicken and flycatcher^43^^,^^44^. We have not found shared syntenic W-linked regions across species that may have been deleted in the ancestor of songbirds compared to the chicken or ostrich genome, as suggested by previous cytological studies^45^.

**Figure 1.**
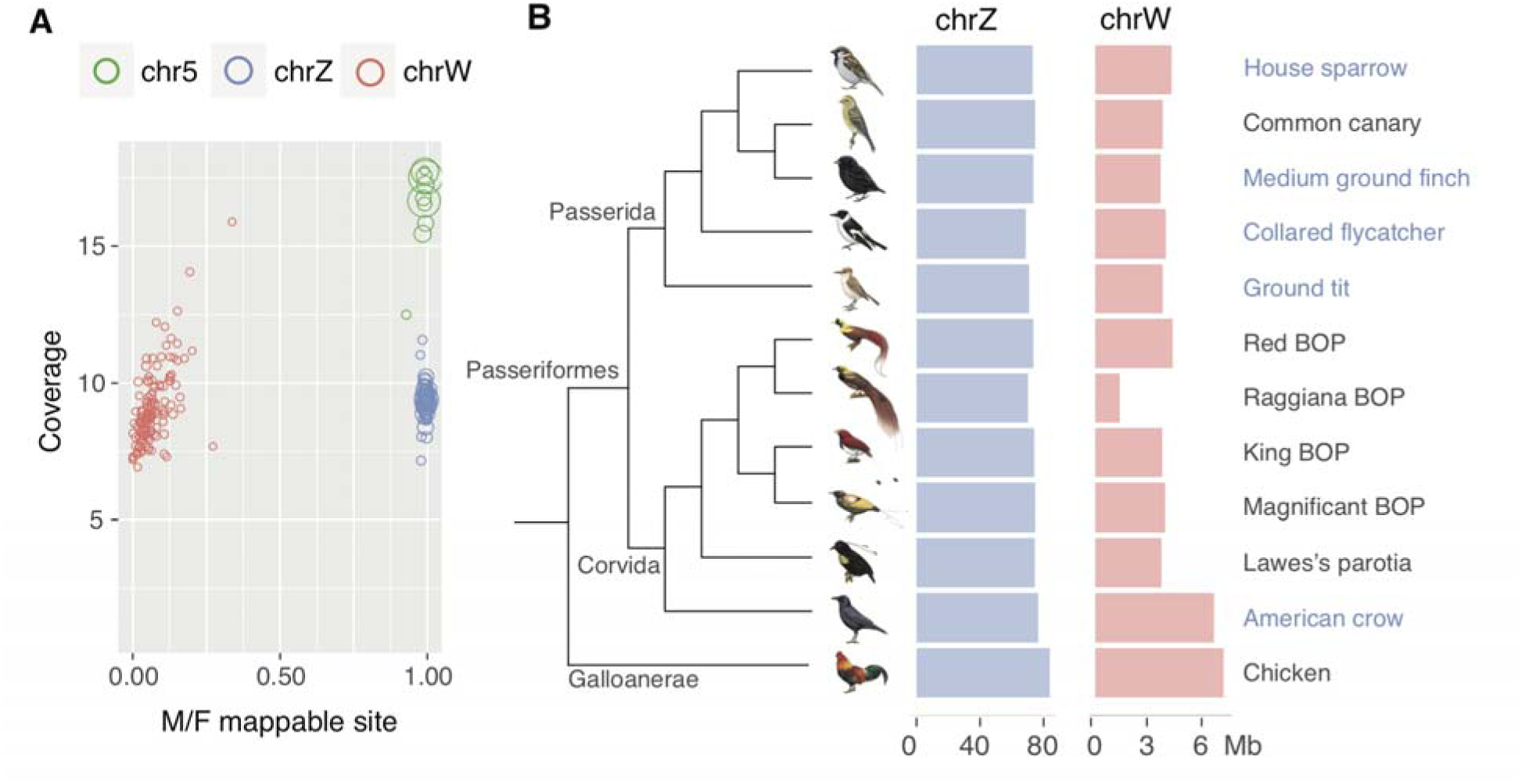
The Z and W chromosomes of different songbirds. **A)** We use medium ground finch as an example to demonstrate our identification and verification of sex-linked sequences. For each scaffold shown as a circle with scaled size to its length, the ratio of nucleotide sites that were mapped by male vs. female genomic reads is plotted against the sequencing depth of this scaffold. Scaffold sequences are clustered separately by their derived chromosomes with W-linked (red circles) and Z-linked (blue circles) sequences showing the expected half the autosome (green) sequencing depth, and W-linked sequences showing almost no mappable sites from male reads. **B)** The lengths of Z and W chromosomes across the studied songbird species. The shorter length of raggiana BOP W chromosome is probably caused by the low sequencing coverage. Species name marked in blue are those that have male reads available for verifying the female-specificity of W-linked sequences.

### Age-dependent genomic impact of evolutionary strata

If recombination was suppressed between sex chromosomes in a stepwise manner, we expect to find a gradient of Z/W sequence divergence levels along the chromosome sequence of the Z chromosome, like what has been reported along the human X chromosome^46^. Previous work showd that the Z chromosomes of the Neognathae have undergone dramatic intrachromosomal rearrangements which resulted in a misleading extant synteny for ordering different evolution strata^20^. By contrast, the Palaeognathae (e.g. emu *Dromaius novaehollandiae* and ostrich *Struthio camelus*) have maintained highly conserved sequence synteny even with reptile species, with over two thirds of their sex-linked regions still recombining as an appropriate approximate of proto-sex chromosomes of all bird species^20^. We first confirmed these patterns by comparative mapping of the nearly chromosome-level assemblies of Z chromosomes of collared flycatcher, hooded crow (*Corvus corone*)^47^ vs. chicken and emu (**Fig. 2A**). This allows us to reconstruct the spanned regions of S0 shared by all birds, and a part of S1 shared by all Neognathae species in the studied songbird genomes by their homology to the emu genome. They are mapped as two continuous regions on the emu Z chromosome, but have become severely reshuffled into dispersed fragments in songbirds (**Fig. 2A, Supplementary Fig. 4-5**). Two recently formed strata (Neoaves S2 and S3) are highly conserved for their synteny across avian species, and each show significantly (*P* < 0.05, Wilcoxon test) different levels of Z/W sequence divergence (**Fig. 2B, Supplementary Fig. 6**), GC3 (GC content at the third codon positions, **Supplementary Fig. 7**) and Z-linked long terminal repeat (LTR) content (**Fig. 2C, Supplementary Fig. 7**) from each other. The drastic change of Z/W sequence similarity allows us to precisely map the boundaries between these two strata. In general, a series of recombination suppression has reshaped the genomic architecture of Z chromosomes in a chronological order. Regions of younger strata exhibit much fewer Z-linked intrachromosomal rearrangements between species, suggesting the reduced selective constraints on gene synteny after recombination was suppressed in the older strata^48^. In particular, GC3 content decreases, while the repeat content increases by the age of stratum. This is probably because of weaker effects of GC-biased gene conversion (gBGC)^49^, and purifying selection against TE insertions^50^ under reduced recombination, both of which have been acting for a longer time within Z-linked regions of older strata. Consistently, a similar pattern has also been found contrasting PAR vs. the remaining Z-linked regions in the collared flycatcher^41^.

**Figure 2.**
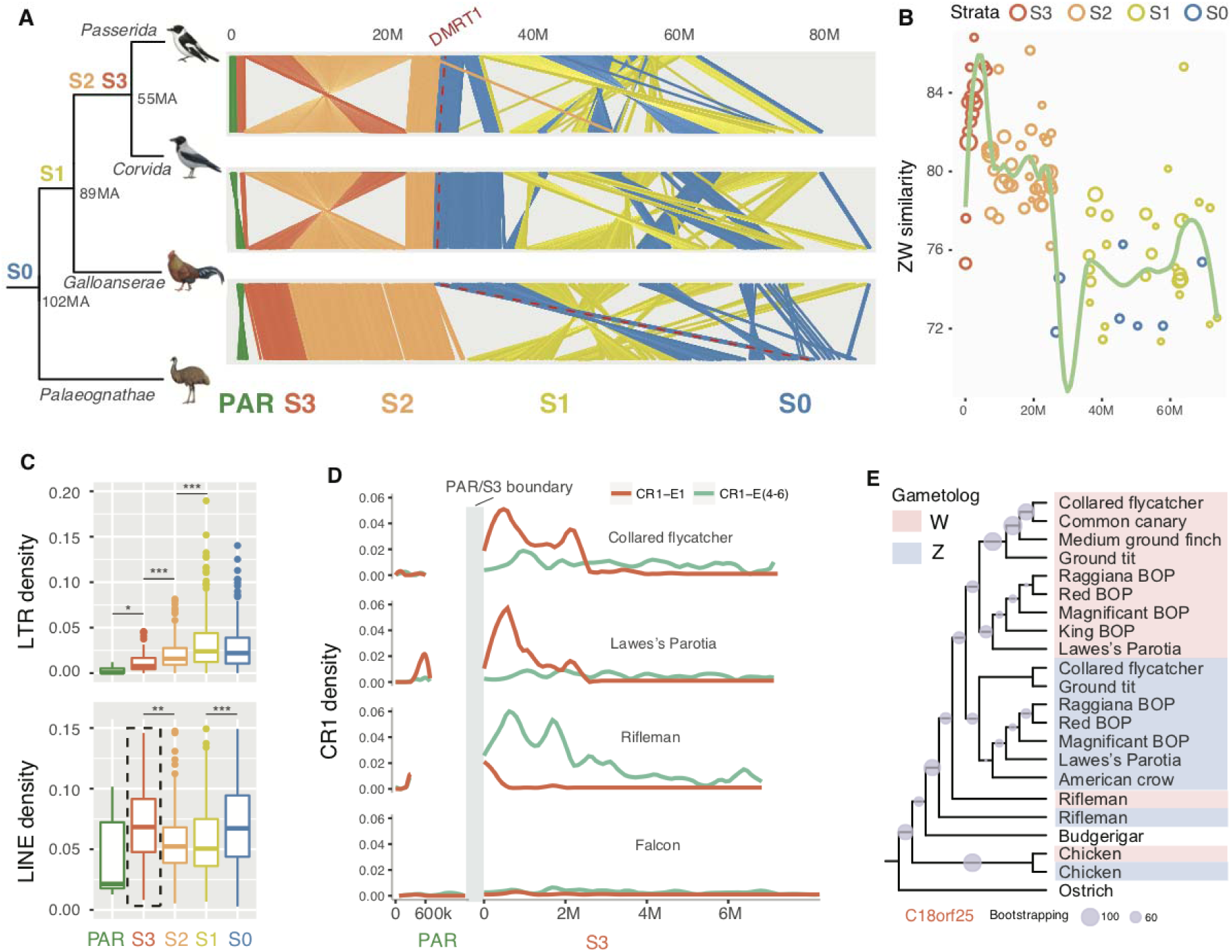
Evolution strata of songbirds. **A)** Genomic synteny of the avian Z chromosomes. Each color represents one evolutionary stratum of songbirds which does not apply to chicken or emu, as they have independent origins of evolutionary strata except for S0. The location of *DMRT1*, the avian male-determining gene is marked by the red dashed line. Generally, the synteny is more conserved in younger strata between species. **B)** We use Lawes’s parotia as an example to demonstrate the pairwise sequence similarity pattern of evolutionary strata. The size of circles is scaled to the length of sequence alignments between Z/W chromosomes. **C)** Transposable elements (LINEs and LTRs) are more strongly enriched in older strata (S0 is the first stratum) except for LINEs at S3. Levels of significance comparing neighboring strata are tested by Wilcoxon test and shown with asterisks: ‘***’: *P*<0.001, ‘**’: *P*<0.01, ‘*’: *P*<0.05. **D)** Lineage-specific burst of CR1-E1 (a subfamily of CR-1 LINEs, red line) at the boundary of the PAR and S3 in songbirds, since their divergence with other passerine species. Other subfamilies of CR1 elements are also plotted with the green line for comparison. **E)** Phylogenetic tree using Z- and W-linked gametolog sequences of the gene *C18orf25* located at S3. Lineages are clustered by chromosomes (red or blue), not by species, suggesting S3 independently formed in rifleman, chicken and the ancestor of songbirds.

### Lineage-specific burst of retrotransposon probably has induced recombination suppression between sex chromosomes

The distribution of long interspersed nuclear elements (LINEs), mainly the retrotransposon chicken repeat 1 (CR1) elements shows an exceptional pattern compared to that of LTR elements (**Fig. 2C**, **Supplementary Fig. 7**) associated with the age of strata. S3 presents a similar proportion of CR1 with S0 that is much higher than the rest of the Z-linked regions. A close examination shows that this is due to the specific accumulation of CR1 elements at the boundary between PAR and S3. Such a burst of CR1, particularly only the CR1-E1 subfamily^51^, extends into about one third of the entire S3 region and is shared by all investigated songbirds but absent in the deep-branching passerine rifleman (*Acanthisitta chloris*) and other Neoaves (**Fig. 2D**, **Supplementary Fig. 8**). The exact boundary sequence is not assembled probably due to such accumulation of CR1-E1, and also previously reported multiple deletions in passerines that removes a gene *DCC* (Deleted in Colorectal Carcinoma) highly conserved across other vertebrates^52^. This gene is responsible for axon guidance for brain midline crossing and has independently been lost in some but not all passerines and Galliformes^53^.

In addition, we find evidence that the burst of CR1-E1 elements coincides with the S3 emergence. Our phylogenetic reconstruction of Z- and W-linked gametolog sequences shows that songbird-derived sequences are always grouped by chromosome instead of species, compared to outgroup birds (**Fig. 2E**, **Supplementary Fig. 9-12**). This indicates that all songbirds share four evolutionary strata, with the latest stratum S3 formed after the divergence between songbirds and the rifleman. The highly conserved synteny between songbird species, and between songbirds and chicken of S3 on the Z chromosome (**Fig. 2A, Supplementary Fig. 5**), suggests that there was no chromosomal inversion on the Z chromosome at S3, and the recent burst of the CR1-E1 subfamily probably contributed to the formation of S3, although we cannot exclude the contribution of possible W-linked chromosomal inversions. Interestingly, other CR1 subfamilies (CR1-E4, CR1-E5, CR1-E6) have an independent burst at the PAR/S3 boundary in rifleman (**Fig. 2D**, **Supplementary Fig. 8** and **Table 3**). Given that this boundary region has been shown to have frequent but different degrees of multiple gene loss in different lineages of birds^52^^,^^53^, it is very likely a hotspot for structural changes (including LINE accumulation) that have recurrently contributed to the independent formation of S3 in many bird species.

### Fast-Z pattern of songbirds implies dynamic evolution of sexual selection

The formation of evolutionary strata has subjected the Z chromosome to male-biased transmission and a reduced effective population size, which are expected to produce faster mutation and evolutionary rates of Z-linked genes^17^. Indeed, we found a larger branch-specific synonymous substitution rate (dS) of Z-linked genes (statistically not significant), but a significantly smaller dS of W-linked genes, compared to that of autosomal genes (*P* < 0.01, Wilcoxon rank sum test, **Supplementary Fig. 13**), as a result of male-driven evolution. The branch-specific evolutionary rates (ω) measured by the ratios of nonsynonymous substitution rate (dN) over dS have significantly (*P* < 0.01, Wilcoxon rank sum test, **Supplementary Fig. 14**) increased for both Z- and W-linked gametologs relative to autosomal genes, indicating a ‘fast-Z’ effect and degeneration of W-linked genes (see below). Previous simulation work and experimental evidence in Galloanserae have suggested that different degrees of sexual selection targeting males will influence the male-mating success, hence the genetic drift effect on the Z chromosome to a different degree^17^^,^^19^. Songbirds, especially BOPs, have been frequently used as textbook demonstrations for sexual selection, though the long-term evolutionary history of sexual selection remained unclear. To reconstruct this, we approximate the intensity of sexual selection targeting males by measuring the fast-Z effect (Z/A value, the ratio of branch-specific ω values of Z-linked genes vs. autosomal genes) in a phylogenetic context (**Fig. 3, Supplementary Table 4**). The varying Z/A values at different lineages suggest a dynamic change of intensity of sexual selection, even among the five BOP species that diverged within the last 15 MY^54^. While the significant (permutation test, *P* < 0.05) fast-Z pattern of the sexually monochromatic American crow (*Corvus brachyrhynchos*) may reflect the sexual selection acting on the ancestral lineage leading to the Corvidae (crows, jays and allies), a lack of such a pattern in the raggiana BOP and magnificent BOP (*Cicinnurus magnificus*) is somewhat unexpected. These species are known for their lekking behaviors^55^, with which very few males dominate almost all females for copulation through outcompeting other males with displays. This produces a strongly biased male-mating success, and direct challenge for maintaining genetic variation in the population (‘the lekking paradox’)^56^. Very few field quantitative studies have been performed on BOP species, and it will be interesting to investigate whether raggiana and magnificent BOPs female individuals may solve the ‘lekking paradox’ by changing their mating preference, and in fact mate with more males than is presumed.

**Figure 3.**
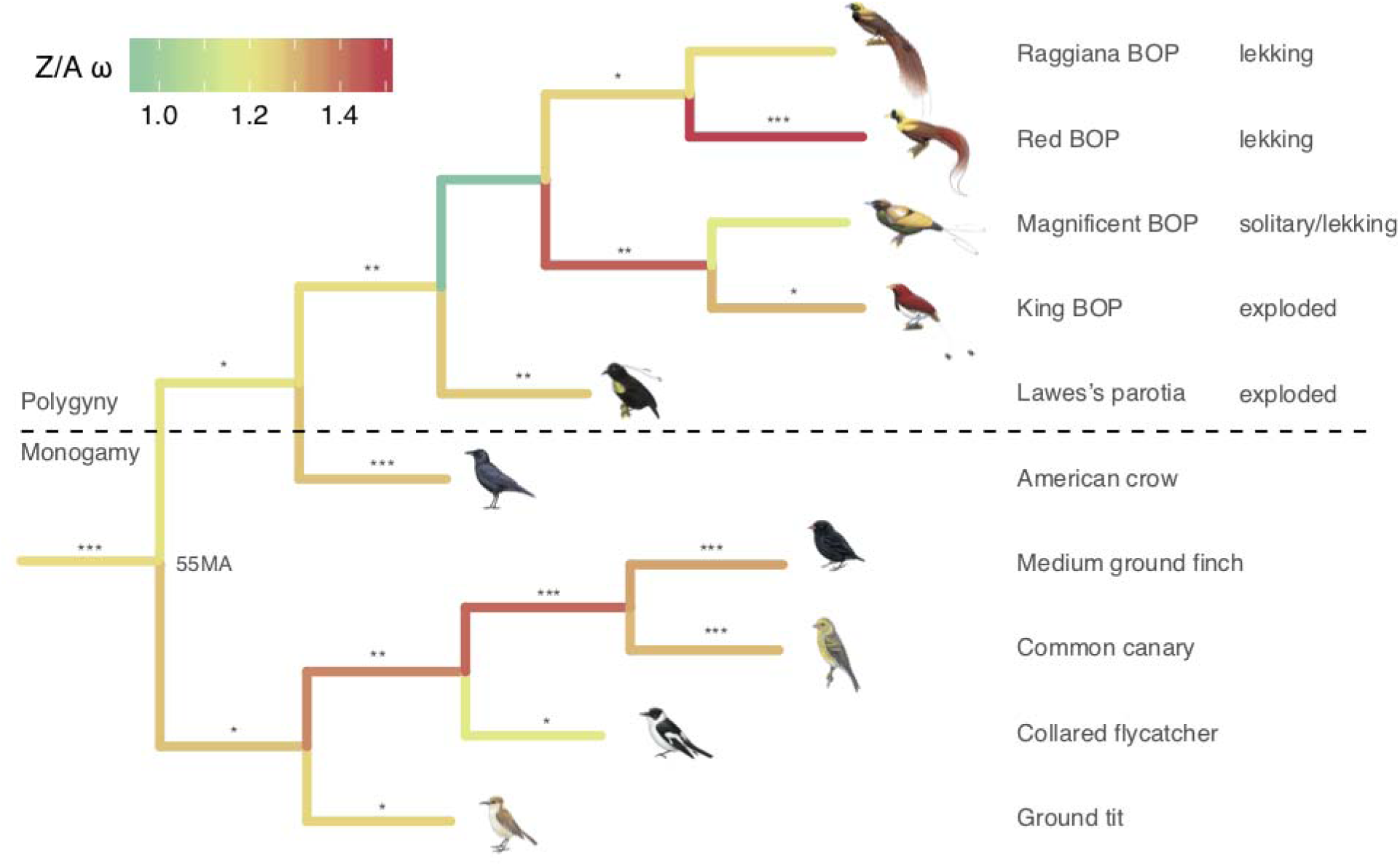
Fast-Z evolution of songbirds. We show the difference of evolutionary rates between Z-linked genes vs. autosomal genes (Z/A value), as a measurement of fast-Z effect throughout the lineages of studied songbird species. The tree length and color is scaled to the Z/A value, with lineages that show a significant (permutation test, *P* < 0.05) fast-Z pattern labelled with asterisks. ‘***’: *P*<0.001, ‘**’: *P*<0.01, ‘*’: *P*<0.05. We also labelled their information mating systems (‘monogamy’ vs. ‘polygamy’), and male display type^54^ (‘lekking’, ‘exploded lekking’, ‘solitary display’).

### Conserved gene content of the songbird W chromosomes

In contrast to the dynamic evolution of Z-linked genes and sequences, W chromosomes of all the studied songbirds have undergone dramatic gene loss but exhibit an unexpected conservation of the retained gene repertoire across species. The numbers of assembled W-linked genes range from 31 in house sparrow (*Passer domesticus*) to 63 in the king BOP (*Cicinnurus regius*), compared to about 600 to 800 Z-linked genes in each species (**Fig. 4A**, **Supplementary Tables 2 and 7**). These numbers are likely an underestimate because genes embedded in highly repetitive regions maybe missing from the current W chromosome assemblies. In general, Corvida species retain more W-linked genes than Passerida species (**Supplementary Table 5**), likely due to their longer generation time thus a lower mutation rate. Most W-linked genes are single-copy without lineage-specific expansion, except for HINT1W (**Supplementary Fig. 15**). Despite the relative rarity of gene duplication in birds compared to mammals^57^, we find one W-linked gene that is a duplicated copy from an autosomal gene *NARF* in American crow (**Supplementary Fig. 16**). This duplicated gene is also present in another Corvida species *Lycocorax pyrrhopterus* (Alexander Suh, personal communication), suggesting that the duplication event likely occurred in the ancestor of Corvida. It will be interesting to see whether this gene shows signatures of female-specific selection, e.g, novel patterns of ovary-specific expression, which drives its fixation on the W. Fifty-seven genes are shared by at least one Corvida and another Passerida species, and 23 genes are shared between at least one songbird species and the chicken^21^. This suggests they were still preserved on the W chromosome before the divergence of passerine or Neognathae species. Interestingly, despite the independent origin of S2 in the chicken and Neoaves^20^, all the chicken W-linked genes but one are also found in passerines, indicating similar underlying evolutionary forces governing their convergent retention since Galloanserae and Neoaves diverged from each other 89 MY ago.

**Figure 4.**
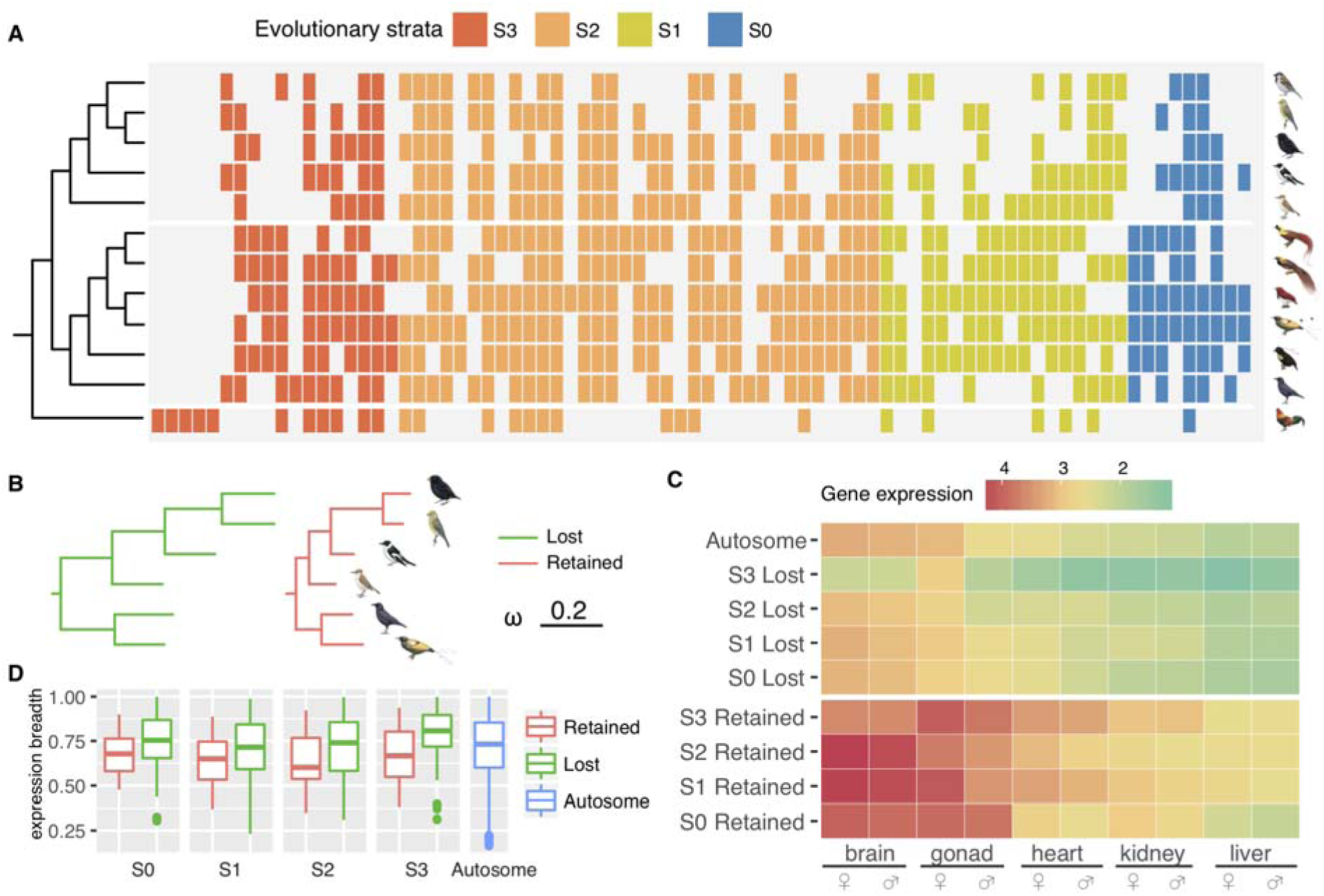
W-linked genes are preserved by purifying selection. **A)** We show the retained W-linked genes of each studied songbird species, as well as those of chicken, with homologous genes aligned vertically. The order of genes follows that of their emu homologs along the Z chromosome. The colors represent the evolutionary strata among songbirds. **B)** The Z-linked genes without W-linked homologs (green, ‘Lost’) evolve faster than those with W-linked homologs retained (red, ‘Retained’), as indicated by their branch lengths scaled to dN/dS ratios. **C)** The Z-linked genes whose W-linked homologs have become lost (upper panel) tend to have a higher expression level (measured by TPM) in their lizard orthologs than those with W-linked homologs retained (lower panel). The genes are divided further by the stratum they reside on, and the expression level is shown by log-transformed medium expression values of each category as color-coded heatmap. **D)** Gene expression tissue specificity in green anole lizard for the homologous avian Z-linked genes.

To examine such forces, we perform gene ontology (GO) analyses on the 79 genes that are present on the W chromosome of at least one songbird species. They are enriched (*P* < 0.01, Fisher’s exact test) for two GO terms of ‘DNA binding’ and ‘transcription factor activity, sequence-specific DNA binding’ (**Supplementary Table 6**). This suggests that similar to the mammalian Y-linked genes^27^, some W-linked genes are retained for their important functions of regulating gene activities elsewhere in the genome. The Z-linked homologs of lost genes evolve significantly (*P* < 0.01, Wilcoxon rank sum test) faster with their ω ratios higher than those of the retained genes on the W chromosome (**Fig. 4B**, **Supplementary Fig. 17**). This shows a different selective pressure acting on these two sets of genes on the proto-sex chromosomes. As this pattern maybe confounded by the ‘faster-Z’ effect of hemizygous Z-linked genes, we further study the autosomal orthologs of these genes in the green anole lizard (*Anolis carolinensis*). We find that the orthologs of retained genes have significantly (*P* < 1.497e-05, Wilcoxon rank sum test, **Fig. 4C**) higher expression levels in all lizard tissues of both males and females, and also a broader tissue expression pattern than those of the lost genes across all the tissues (**Fig. 4D**). The patterns are also consistent among the four songbird evolutionary strata. These results are robust if we use emu to infer the ancestral expression pattern (**Supplementary Fig. 18**), whose sex chromosomes are largely a PAR. Consistent with the result of collared flycatcher^10^, we find no evidence that female-specific selection may prevent the gene loss, or drive certain genes to undergo positive selection on the songbird W chromosomes. We found no excess of ovary-biased lizard orthologs among those of the retained W-linked genes: only 6 out of 72 (8.3%) are ovary-biased while the genome-wide proportion is about 20%.

### Comparing gene loss between avian W chromosomes and mammalian Y chromosomes

Overall, 4.6% to 9.2% of single-copy W-linked genes per songbird species (**Fig. 4A**), compared to 1.6% to 3.0% single-copy Y-linked genes per mammalian species^27^ have been retained for their essential or sex-specific functions. A seemingly high retention ratio of W-linked genes in birds can be partially attributed to the generally much lower mutation rate of W chromosome relative to Y chromosome by male-driven evolution effect (**Supplementary Fig. 13**), assuming a similar generation time between mammals and birds. In addition, a much more frequent and stronger sex-specific selection acting on the Y chromosome than on the W chromosome, sometimes driving the massive expansion of Y-linked gene copies with male-related function^24^, probably also has contributed to a faster rate of Y chromosome gene loss by the hitchhiking effect^24^^,^^26^. To scrutinize the tempo of gene loss throughout the evolution of songbird sex chromosomes, we conservatively reconstructed the numbers of retained W-linked gametologs at each phylogenetic node of the avian tree (**Fig. 5A, Supplementary Table 8**), by identifying the genes present on any of the studied avian W chromosomes. We find that within each stratum, the percentage of gene loss is always much larger at an earlier evolutionary time point than in the recent ones, and this is consistent between birds and mammals (**Fig. 5B**). Thus the majority of gene loss has probably occurred during the early stage of W or Y chromosome evolution, and the rate of gene loss dramatically decreases by the less retained genes. Although convergent gene loss may cause an overestimation of lost genes at more ancestral time points (e.g., in S0 region), this probably has little influence on our estimate in the most recent songbird-specific stratum S3 which has already lost 69.8% of the W-gametologs within 50 MY. We also find that the retained genes of songbird W chromosomes are often close to each other (**Supplementary Fig.19**), suggesting large sequence deletions have had an important contribution to drastic gene loss.

**Figure 5.**
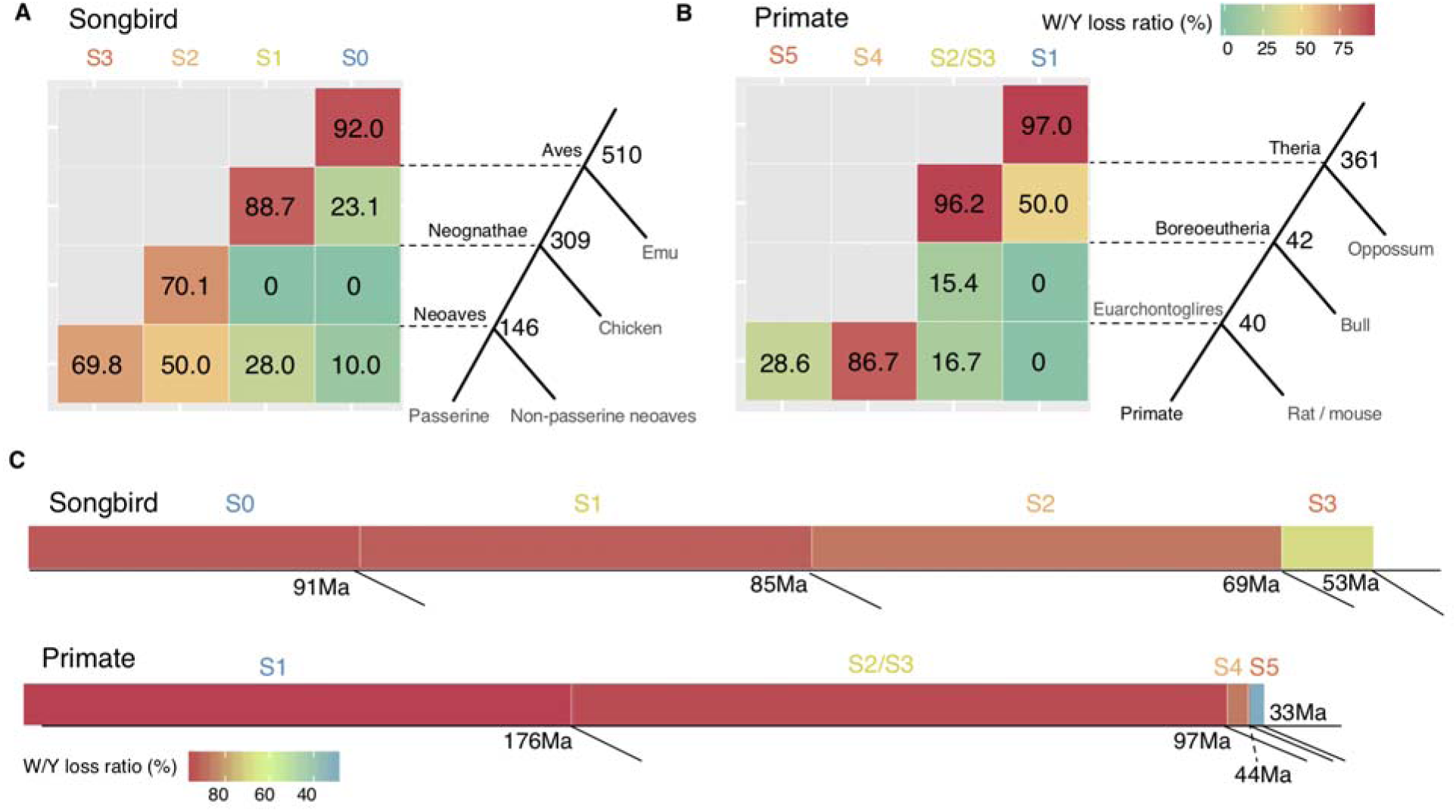
Comparison of gene loss between W chromosomes of songbirds and Y chromosomes of primates. **A)** We show the percentage of gene loss, and ancestral gene number for each evolutionary stratum at each phylogenetic node. **B)** Similar analyses for the Y-linked gametologs of primates based on the data of ^21^, with S1 as the first stratum of eutherian mammals. **C)** We show the length of songbird W or primate Y chromosomes scaled to the ancestral gene number of each evolutionary stratum, with the color scaled to the overall percentage of gene loss. The ages of evolutionary strata are indicated by the number (in millions of years) at the nodes below the bars. As eutherian mammals have much larger ancestral evolutionary strata than those of birds, they probably suffer a more severe gene loss on the Y chromosome.

The decrease of gene loss rate on Y/W chromosomes through evolutionary time can be explained by a weaker Hill-Robertson effect that the less retained genes can induce, which has been previously shown in a simulation study^26^. Thus, the ancestral gene number of older evolutionary strata which would have undergone more serious gene loss must have a larger influence on the extant number of retained genes. A lower rate of retained mammalian Y-linked genes relative to avian W-linked genes, besides the cause of male-driven evolution, can be also attributed to the fact that the first two or three mammalian evolutionary strata emerged before the divergence of eutherians which together account for over 93.2% of the entire gene content of ancestral Y chromosome, while those of birds only account for 53.3% of the entire ancestral W-linked gene content (**Fig. 5C**).

## Discussion

The evolution of sex chromosomes is usually, but not always (e.g., in frogs^58^, ratite birds^20^ and pythons^59^), marked with episodes of recombination suppression that eventually restricts the recombining region to one or two small PARs at the end of the sex chromosomes. The resulting patterns of evolutionary strata have been widely reported in many animal and plant species, with the responsible formation mechanism presumed to be chromosomal inversions^60^. Indeed, footprints of inversions in the latest two strata between mammalian X and Y chromosomes have been found by examining the synteny order between X/Y, and particularly the X-linked boundary sequence that has been disrupted into two dispersed sequences on the Y chromosome^61^. In songbirds, we show here that four evolutionary strata were formed before their species radiation. Importantly, we provide evidence suggesting an alternative scenario of recombination suppression. We find in the latest stratum S3, a genome-wide burst of a certain transposable element (TE) led to its specific accumulation at the mutation/insertion hotspot near the PAR boundary of the sex chromosomes (**Fig. 2, Supplementary Table 3**), which probably further mediated recombination suppression. It has been reported with some exceptions, in many species that local recombination rates and abundance of TE elements generally have a negative association, with their causal relationship difficult to be disentangled^50^. In the case of the songbird S3 region, several patterns suggest that TE accumulation might be the cause rather than the result of recombination suppression. First, in several species (e.g., Lawes’s parotia (*Parotia lawesi*) and the king BOP, **Fig. 2D**, **Supplementary Fig. 8**), the responsible CR1-E family is also enriched close to the PAR boundary, where normal levels of recombination can be expected. Second, the abundance of CR1-E gradually decreases further away from the boundary, which also has undergone genomic deletions or rearrangements independently in other bird species^52^^,^^53^, thus is likely to be a mutation hotspot. Third, only CR1-E repeats, but not any other type of CR1 or repeat families, are enriched at the PAR/S3 boundary. These results are not likely to be expected if S3 stratum formation occurred before the CR1-E accumulation, which predicts a uniformly distributed accumulation of various kinds of repeat elements (e.g., LTR elements, **Supplementary Fig. 20**) by Hill-Robertson interference effects that would not extend into the PAR. Indeed, our comparative analyses between species suggest no Z-linked inversion in S3 (**Fig. 2A, Supplementary Fig. 5**), though we cannot exclude the possibility of a W-linked inversion that may have instead contributed to the formation of S3. The verification of the latter requires future improvements of genome assembly using for example, PacBio/Nanopore sequencing technology to assemble the highly repetitive W-linked sequence.

Under our proposed scenario, TEs probably reduced the recombination rate in PAR through, for example, changing the chromatin structure or disrupting recombination hotspots^50^. The CR1-E retrotransposon accumulation occurred at the mutation hotspot located at PAR/S3 boundary despite the selection against its deleterious effects of disrupting gene functions. This has resulted in partial or complete deletion of several genes in songbirds^52^^,^^53^. However, the consequential reduction of recombination rate can provide the selective advantage of accelerating the fixation of pre-existing sexually antagonistic (SA) alleles in PAR through sex-biased transmission; or subjecting the PAR for the ‘fast-Z’ evolution by male-driven evolution effect (**Fig. 3**) and increasing its exposure for male-biased selection, so that novel SA alleles may more frequently emerge and become fixed. The latter has been implicated by the recent findings in songbirds that male-specific trait genes, for example those related to sperm morphology^62^ or plumage colors^63^, which have recently become diverged within or between populations, are enriched on the Z chromosome. In addition, TE accumulation is likely to increase the chance of chromosome inversions through ectopic recombination, or by reducing the selective constraints on gene synteny^48^^,^^50^. The latter is supported by our result that older evolutionary strata have undergone more Z-linked genomic rearrangements between songbird species than the younger ones (**Fig. 2A**), which creates a positive feedback once the recombination suppression was initiated. This provides a mechanistic explanation for the more frequent fixation of Z-linked inversions found among passerines.

While the Z chromosome is predicted to accumulate dominant male-beneficial mutations, the W chromosome is expected to accumulate female-beneficial mutations responding to female-specific transmission^64^. Here we find that more genes to have been retained on the W chromosomes of songbirds than that of chicken (**Fig. 4**). However, both previous works on the chicken and collared flycatcher^21^^,^^30^, as well as our present study, have not found evidence for such ‘feminization’ of W chromosome. This is in contrast to multiple reported cases of ‘masculinization’ of the Y chromosome in the ancient mammalian Y chromosome systems^24^ or the recently-evolved Y chromosome of *Drosophila miranda*^28^. Genes specifically expressed in the male germline have either massively amplified their copy numbers, or upregulated their expression levels on these evolving Y chromosomes. Such a difference can be explained by the fact that regardless of the sex chromosome type, sexual selection is often targeting males in most species, thus the Z/Y chromosomes are more frequently influenced than the W/X chromosome due to their male-biased transmission, although X chromosomes are nevertheless expected to accumulate recessive male-beneficial alleles^9^. The convergently evolved pattern shared between the mammalian Y and avian W chromosomes is largely attributed to the essential genes that have important regulatory functions and are preferentially retained over the long period of recombination suppression (**Fig. 4**)^21^^,^^27^. However, previous transcriptome comparison of chicken breeds selected for egg-laying vs. fighting, i.e. female-specific vs. male-specific traits, has found that most W-linked genes are upregulated in the former^64^. Few high-quality avian W chromosome sequences are available except for that of chicken^21^^,^^65^. Songbirds provide a rich resource with many species having a reversed direction of sexual selection and ornamented females^66^. Application of long-read sequencing technology in future will help to elucidate the role of the W chromosome in sexual selection and speciation of birds^67^.

## Materials and Methods

### Genome assembly and annotation

Genomic DNA were extracted from fresh female tissue samples of from BOP species *Cicinnurus regius* (ANWC B24969), *Cicinnurus magnificus* (ANWC B27061), *Paradisaea rubra* (YPM84686), *Parotia lawesii* (ANWC B26535), using Thermo Scientific™ KingFisher™ Duo Prime purification system. Paired-end and mate pair libraries for these samples were prepared by SciLifeLab in Stockholm, Sweden. All libraries were sequenced on Illumina HiSeq 2500 or Hiseq X v4 at SciLifeLab in Stockholm, Sweden. For *Paradisaea raggiana* (USNM638608), genomic DNA were extracted using EZNA SQ Tissue DNA Kit. One paired-end and one mate-pair libraries were sequenced on Illumina HiSeq 2000. We also used published female genomes of *Corvus brachyrhynchos*, *Serinus canaria, Passer domesticus, Geospiza fortis, Ficedula albicollis, Pseudopodoces humilis* for analysis in this work (**Supplementary Table 9**)^30^^,^^36^^-^^39^. The BOP genomes were assembled using ALLPATHS-LG^68^ with ‘HAPLOIDIFY=True’. For *P. raggiana*, due to the lack of overlapping paired-end reads, SOAPdenovo2^69^ was used instead. Gene models were annotated using the MAKER pipeline in two rounds^70^. The reference protein sequences of zebra finch, great tit, hooded crow, American crow, collared flycatcher and chicken were downloaded from NCBI RefSeq (**Supplementary Table 9**). Using the reference protein sequences and chicken HMM (Hidden Markov Models), an initial set of gene models was obtained by using MAKER, and those models were taken for SNAP model training^71^. In addition, 3000 gene models were selected for Augustus training^72^. The trained gene models and the protein sequences were taken as input for MAKER in the second run. To annotate repeats, first we used RepeatModeler^73^ to identify and classify repeat elements for each species, including the published genomes^74^. Then we combined each individual library with an avian repeat library and further manually curated BOP repeat consensus sequences, followed by annotation repeats using RepeatMasker (v1.7)^73^.

### Identification of sex-linked sequences

We used the sequence of the great tit Z chromosome as reference to search for homologous Z-linked sequences in the studied species, as it has the best assembly quality (contig N50: 133kb, scaffold N50: 7.7Mb) among the published genomes of songbirds^6^. We used nucmer from MUMMer package (v4.0)^75^ for genome-wide pairwise sequence alignment. For any scaffold larger than 10 kb that has more than 60% of the sequence aligned to great tit chrZ, it was identified as candidate Z-linked sequences. They were further inspected by comparing the female sequencing coverage to that of autosomes. To do so, the raw female reads were mapped to the genome references by using bwa^76^ then the mean sequencing depth of every 50-kb window was calculated. Sequencing depth for single sites was counted using ‘samtools depth’ before calculating window-based coverage. Any site with low mapping quality (less than 60) and very high coverage (3 times larger than average) was excluded.

To identify the W-linked scaffolds, first we identified the scaffolds that show half-coverage relative to that of autosomes. We plotted the distribution of coverage of all scaffolds to decide the cutoff of ‘half-coverage’ (**Supplementary Fig.1**). Those half-coverage scaffolds were expected to be either Z-linked or W-linked. To distinguish between the Z and W, we aligned the half-coverage scaffolds to the Z chromosome from the hooded crow, a closely related species to BOP, whose genome is derived from a male without contamination of W-linked sequences. We used nucmer for sequence alignment and only kept 1-to-1 best alignments. We then calculated the proportion of sequences of each scaffold that was aligned to the hooded crow chrZ, and decided a cutoff to separate the Z and W based on the distribution of the proportion of Z-linked alignment. We further excluded the candidate W-linked scaffolds that over 10% of the sequences were aligned to hooded crow autosomes, or a larger portion of sequences aligned to the Z chromosome than the W. Finally, only the scaffolds that were larger than 50 kb were kept. We also retrieved additional Z-linked scaffolds that were absent in the results from the homology-based approach, likely due to the missing Z-linked sequences in the great tit Z chromosome assembly. For ground tit, medium ground finch, house sparrow and collared flycatcher in which both male and female sequencing reads are available, the W-linked sequences were further verified by mapping the male reads. Specifically, for every scaffold, the number of nucleotide sites that were covered by male and female sequencing data were counted respectively as Nm and Nf, and the ratios of Nm to Nf were calculated. W-linked scaffolds were expected to have Nm/Nf ratios close to zero and one for autosomal or Z-linked scaffolds. For PARs, we used the known sequences of the zebra finch^77^ and collared flycatcher^41^ to search for the homologous sequences in other species using nucmer. The female sequencing depths of those candidate PARs were compared to autosomes and required to be similar.

### Demarcation of evolutionary strata

We ordered and oriented the identified Z-linked scaffolds into one pseudo-chromosomal sequence (pseudo-chrZ) based on their alignments against the chromosomal assembly of great tit. Hooded crow has only 15 Z-linked scaffolds and 10 out of them are larger than 1 Mb^47^, thus was used as a representative Corvida species for comparison on the Z chromosome. We determined the relative order and orientation of the scaffolds according to their alignment on the great tit Z chromosome. Similarly, for BOP species, we created pseudo-chromosome Z using great tit as guiding reference. The pseudo-Z chromosome of emu was built using ostrich Z chromosome^78^ ^79^as reference. We used nucmer for pairwise alignment of the Z or pseudo-Z chromosomes. Alignments short than 2kb were excluded.

The W-linked scaffolds were then aligned to the pseudo-chrZ using lastz^80^, after masking repetitive sequences. Sequence similarity of the alignments between the chrZ and chrW was calculated by the script pslScore from UCSC Genome Browser (https://genome.ucsc.edu/). Individual alignments that had sequence similarity lower than 60 or higher than 96, or alignment length shorter than 65 were removed. After that, we concatenated alignments within non-overlapping sliding windows of 100 kb, and calculated sequence similarity for the concatenated alignments. When the length of concatenated alignments was shorter than 2 kb within a 100-kb window, the window was excluded from further analyses. The window-based sequence similarity was then plotted along the pseudo-chrZ. The shift of sequence similarity was used to demarcate the boundaries of S3/S2 and S2/S1. Since very few W-linked sequences have been assembled for the most ancient stratum S0, we mapped its reshuffled fragments in songbirds based on their homology with the emu S0. Our previous study showed emu has a recent species-specific stratum (S1) while the first stratum (S0) is ancient and shared by all birds^20^. This allows for the demarcation of S1 and S0 by detecting their differential degree of ZW differentiation. Specifically, by using relatively relaxed mapping criteria (bwa mem) to map female sequencing reads, only S0 showed reduced coverage relative to autosomes or PAR (**Supplementary Fig. 4**), while S1 showed reduced coverage when stringent mapping was applied (bwa sampe –a 900 –n 1 –N 0 –o 10000).

To scrutinize the accumulated LINE (mostly CR1) elements at the PAR/S3 boundary, we first divided them into each subfamily (approximated by each repeat consensus sequence) according to the RepeatMasker annotation. Among all subtypes, CR1-E1 is usually ranked with the highest or second highest number at the S3 region across all songbird species. Other high-ranking subtypes included CR1-E3, CR1-E5, CR1-E4, CR1-E6, CR1-J2 and CR1-Y2. Then we plotted each subtype’s abundance with a 100 kb non-overlapping window along the Z chromosome, in all the studied songbirds, as well as outgroup species rifleman and falcon^81^, to identify the burst of CR1-E1.

### Sex-linked gene analyses

After removing LTR-derived genes, we used BLAT^82^ to align the annotated coding sequences of W-linked genes to the Z chromosome to search for their gametologous pairs. Then we produced pairwise gametolog alignments using MUSCLE,^83^ and then manually inspected the alignments to remove genes with short or ambiguous alignments. For species other than the BOPs, gene models of the W chromosomes were directly retrieved from the RefSeq genome annotation, with some of them subjected to manual inspections. To determine the orthologous relationship among the studied species, we first extracted the sequence of the longest protein of each gene. Those protein sequences were subjected to all-vs-all BLAST search that was implemented through the program proteinortho^84^. BLAST hits with identity lower than 50% or alignment coverage lower than 50% were removed. We also took gene synteny information into account when grouping orthologous genes. Besides the twelve female genomes for which we studied the sex chromosomes, we also included high-quality genomes of great tit, hooded crow and ostrich (**Supplementary Table 9**). We retained those orthologous groups if they contained sequences of at least ten species.

To estimate the substitution rates of coding sequences, first we performed multiple sequence alignments for orthologous genes. We used the guidance2 pipeline (http://guidance.tau.ac.il/ver2/source.php) which employs PRANK to align sequences of codons. To filter low-quality sites in the alignments, we ran trimal (http://trimal.cgenomics.org/) to further filter the ambiguous alignments with ‘-gt 0.8’. The phylogeny of the birds was extracted from Jetz et al. (2012)^85^. We used codeml from the PAML package^86^ to estimate the synonymous substitution rates (dS) and non-synonymous substitution rates (dN). To estimate chromosome-wide dN and dS, the sums of synonymous or nonsynonymous substitutions were divided by those of the number of synonymous or nonsynonymous sites, as applied in Wright et al.^19^. Individual genes with abnormal dN (higher than 0.1, in total 179 genes) or dS (higher than 0.8, in total 135 genes) out of 111,748 orthologous gene groups were removed. Confidence intervals were calculated by 100 bootstraps. The GC content of the third position codons (GC3) was calculated using codonW^87^ for the longest isoform of each gene. Chromosome-wide dN/dS (ω) was calculated using the ratios of chromosome-wide dN to chromosome-wise dS. The fast-Z effect is measured by Z/A value, the ratio of ω values of Z-linked genes to autosomal genes, and we calculated the Z/A value for every terminal branch and internal branch. To determine whether the difference of ω between Z-linked and autosomal genes wasis significant, we performed permutation test by resampling 1000 times. The genes of chromosome 4 and chromosome 5 were used to represent autosomal genes as the sizes of those two chromosomes are similar to the Z chromosome.

For each gametologous pair, we grouped together Z-linked genes and assembled W-linked genes and performed multiple sequence alignment. The same guidance2 pipeline was used as in sequence divergence analysis. For S3 genes, we also included rifleman^36^. We used IQTREE^88^ to construct maximum likelihood phylogenetic trees. The best substitution model was automatically selected in by IQ-TREE. We ran 100 bootstraps to evaluate the confidence levels of phylogenies. Ostrich was used as the outgroup to root the tree. The gene ontology (GO) term annotations for both gametolog-pairs genes (list) and entire Z-linked genes (background) of chicken were analyzed using DAVID 6.8^89^. GO term enrichment was analyzed by comparing the number of appearance of GO terms of ‘list’ gene versus ‘background’ gene.

### Gene loss analysis

We identified a total of 673 Z-linked orthologous genes that are shared between chicken and emu as the putative ancestral genes on the proto-sex chromosomes of birds. For the gene cluster that was lost in chicken at the DCC locus of S3, an ancestral gene content was inferred based on Fig. 3 of Patthey et al.^53^. They were then grouped into four evolutionary strata according to the strata annotation of songbird Z chromosomes. At each node of the avian phylogenetic tree, we calculated the ratio of the number of lost genes to the number of ancestral genes at that node. For the nodes leading to Passerida and Corvida, if there were at least one species retaining a W-linked gene, we inferred that this gene was present in their ancestor. Similarly, we defined the presence of ancestral genes in Passeriformes, Neoaves and Neognathae using other published avian W-linked gene information^20^^,^^21^.

### Gene expression analysis

We downloaded the raw RNA-seq reads of green anole (brain, gonad, liver, heart and kidney) and emu (brain, gonad and spleen) from SRA (**Supplementary Table 9**). In addition, we collected transcriptomes of adult emu kidneys of both sexes. We used the RSEM pipeline^90^ to quantify the gene expression levels. We used STAR^91^ to map raw reads to the transcriptomes which was constructed based on gene annotations. The expression levels were estimated at the gene level, in the form of TPM (Transcripts Per Million). The mean TPM value of biological replicates was calculate for each gene. Tissue specificity of gene expression was estimated by calculating tau^92^.

### Data availability

Genome sequencing and RNA-seq data, and genome assemblies generated in this study have been deposited in the NCBI SRA under PRJNA491255.

## Acknowledgements

We thank Australian National Wildlife Collection, CSIRO Sustainable Ecosystems (Leo Joseph), Museum Victoria, Australia (Joanna Sumner), Division of Vertebrate Zoology Yale University, Peabody Museum of Natural History (Kristof Zyskowski) for tissue samples; and Edwin Scholes for discussions on birds-of-paradise. We also acknowledge the support from Science for Life Laboratory, the National Genomics Infrastructure (NGI), Uppmax. L.X. is supported by the uni:doc PhD. fellowship from University of Vienna. M.I. is supported by the Swedish Research Council (grant number 621-2014-5113). A.S. is supported by the Swedish Research Council (grant number 2016-05139) and thanks Muhammad Bilal for help with repeat annotations. Q.Z. is supported by National Natural Science Foundation of China (31722050, 31671319), the Thousand Talents Plan, the Fundamental Research Funds for the Central Universities, and start-up funds from Zhejiang University. The computational analyses were performed on CUBE cluster from Dept. of Computational System Biology of University of Vienna and Vienna Scientific Cluster.

